# Abnormal eNK cells contribute to endometrial fibrosis in intrauterine adhesions patients

**DOI:** 10.1101/2024.12.01.626271

**Authors:** Zhenhua Zhou, Qiao Weng, Dan Liu, Simin Yao, Xueqin Zou, Huiyan Wang, Zihan Zhou, Hui Zhu, Xier Zhang, Ling Chen, Xiwen Zhang, Guangfeng Zhao, Yali Hu

## Abstract

Endometrial natural killer cells (eNK) at the proliferative phase are one of the major lymphocytes in the endometrium, but their phenotype and function remain elucidated. Here, we demonstrated that eNK cells had three subtypes and were different from peripheral blood NK (pbNK) and decidual NK (dNK) cells. They played an important role in maintaining the homeostasis of the endometrium through preventing endometrial stromal cells (ESCs) from the differentiation into myofibroblasts physiologically. However, in the fibrotic endometria of patients with intrauterine adhesions (IUA), eNK cells were significantly decreased and highly negatively correlated with an increase in myofibroblasts. In vitro experiments showed that eNK cells could inhibit the differentiation of ESCs into myofibroblasts and promote the dedifferentiation of myofibroblasts, in which the main effector molecules from eNK cells was Prostaglandin D2 (PGD2). PGD2 downregulated the expression of ZNF521 to decrease profibrotic protein synthesis. Furthermore, we confirmed the anti-endometrial fibrosis effect of eNK cells and its mechanism in an IUA-like murine model. These findings reveal an important role of eNK cells in endometrial homeostasis and provide potential therapeutic approaches for IUA patients.

## Introduction

Intrauterine adhesion (IUA) is the most common cause of female uterine infertility [1–3]. It refers to the endometrial basalis injury mainly caused by trauma [4], in which the endometrial regeneration is impaired and replaced by a large amount of fibrous tissue, resulting in partial or complete occlusion of the uterine cavity [3]. Although hysteroscopic adhesiolysis, as the preferred treatment for IUA, can restore the uterine cavity morphology of patients with severe IUA, the re-adhesion is over 60% [1, 5]. Most importantly, endometrial fibrosis is difficult to be reversed resulting in a refractory infertility.

Endometrial fibrosis is characterized by the differentiation of large number of endometrial stromal cells (ESCs) into myofibroblasts, which are marked by the expression of α-smooth muscle action (α-SMA), and the production of a large amount of extracellular matrix (ECM), especially collagen [6–8]. During tissue injury, myofibroblasts contribute to wound healing and are then immediately removed upon completion of repair [8]. If they are persistently increased, extensive deposition of ECM, endometrial stiffness and distortion of the uterine cavity will occur [7–9]. Therefore, the dynamic balance of formation and dedifferentiation of myofibroblasts is crucial for maintaining the physiological function of the endometrium. Hence, in-depth exploration of the underlying mechanism of endometrial fibrosis is of great significance for more effective guidance of endometrial functional reconstruction.

Natural killer (NK) cells are an important lymphocyte subset of innate immunity and play an important role in immune surveillance, killing and timely elimination of mutated or abnormal cells to maintain tissue homeostasis [10–14]. NK activation depends on ligand-receptor interaction and is an outcome of the balance between activation and inhibition signals [15]. Uterine NK cells include endometrial NK (eNK) cells and decidual NK (dNK) cells depending on the phase of the menstrual cycle [16]. dNK cells comprise up to 70% of leukocytes in first trimester of pregnancy and play an important role in mediating maternal-fetal immune tolerance, promoting placental/fetal development, eliminating microbial infection, and limiting the spread of placental inflammation [17–21]. Therefore, without properly functioning dNK cells, there is no normal pregnancy. However, so far, the physiological functions of eNK cells in the proliferative endometrium remain unclear.

Here, we demonstrated that eNK cells had three subtypes and were different from peripheral blood NK (pbNK) and dNK cells. They played an important role in maintaining the homeostasis of the endometria through preventing ESCs from the differentiation into myofibroblasts physiologically and in the endometria of IUA patients, eNK cells and their secretion of PGD2 were significantly decreased, resulting in increased formation and decreased dedifferentiation of myofibroblasts to contribute endometrial fibrosis, which was related to up-expression of ZNF521.

## Results

### Decreased eNK cells is related to increased myofibroblasts in the endometria of IUA patients

To decipher the endometrial cell atlas in IUA patients, we used single cell RNA sequencing (scRNA-seq) data of endometria in proliferative phase including normal controls (n = 3) and severe IUA patients (n = 3) [22]. After quality filtration, we obtained 57845 cells composed of 28381 cells from normal endometria and 29464 cells from IUA endometria, and then the total cells were cataloged into 14 distinct clusters based on Single R annotation and known marker genes (Fig. 1A, B, Supplementary Fig. 1A). Compared with normal controls, the proportion of fibroblasts changed obviously, among which the proportion of proliferative fibroblasts was decreased (6.62% vs 8.99%), while the proportion of ACTA2+ fibroblasts, highly expressed ACTA2 and PDGFRB, was increased (5.53% vs 4.33%) (Fig. 1C). As shown by immunohistochemical staining, in normal endometrial tissues, α-SMA positive cells were mainly located around blood vessels, namely perivascular cells (Supplementary Fig. 1C). However, in IUA endometria, they were not only around blood vessels, but also distributed in the stromal compartment, suggesting an increased myofibroblasts in IUA endometria. Masson staining showed a large amount of collagen deposition, which was consistent with endometrial fibrosis (Supplementary Fig. 1B, C). Concurrently, the proportion of immune cells was also disrupted, among which eNK cells unexpectedly declined most in endometria of IUA patients (15.74% vs 25.33% of immune cells) (Supplementary Fig. 1D). To verify the changes of eNK cells in IUA analyzed by scRNA-seq, we performed the flow cytometry, immunohistochemistry and immunofluorescence staining of endometria. The results showed that the proportion and number of eNK cells reduced significantly in IUA patients (Fig. 1D-G). What’s more, a highly negative correlation between the proportion of eNK cells and ACTA2+ fibroblasts was found (Fig. 1H). And CellPhoneDB analysis of eNK cells also revealed a significant interaction between NK cells and ACTA2+ fibroblasts (Supplementary Fig. 1E), indicating that eNK cells might be involved in myofibroblasts differentiation and endometrial fibrosis.

**Fig. 1.**
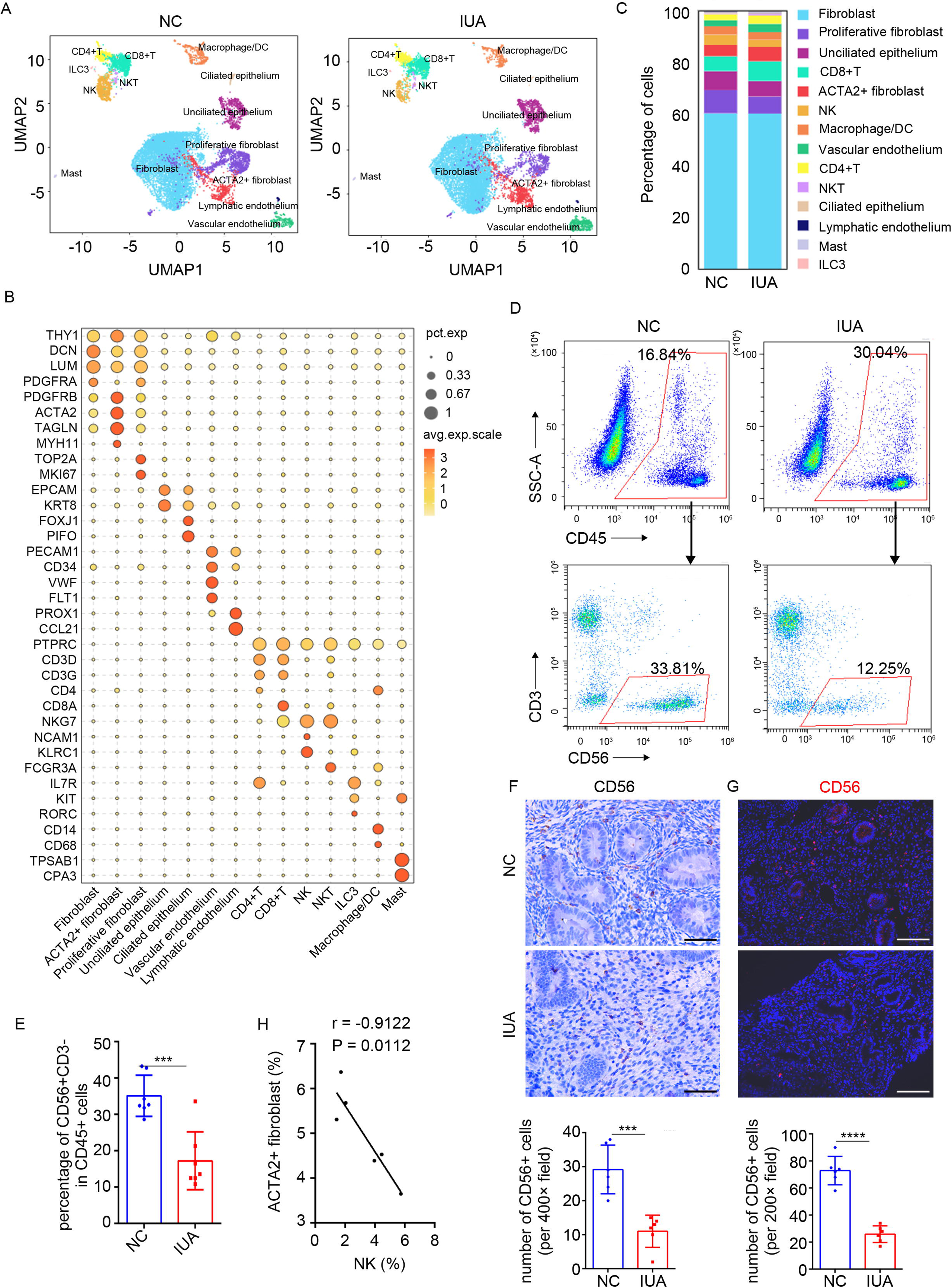
eNK cells are decreased in endometria of IUA patients. (A) The UMAP plot showing the distribution of 14 major cell clusters in normal and IUA endometria. (B) Bubble plot showing the expression of marker genes for each cell cluster. (C) The proportion of each cell type in normal controls (n = 3) and IUA patients (n = 3) based on scRNA-seq. (D) Flow cytometric analysis of proportion of eNK cells (gated by FVS510-CD45+CD3-CD56+) in endometria of normal controls (n = 7) and IUA patients (n = 7). (E) Statistical plot of the proportion of eNK cells analyzed by flow cytometry. (F) Immunohistochemical of CD56 in endometria of normal controls (n = 6) and IUA patients (n = 6). Scale bars: 50 μm. (G) Immunofluorescence staining of CD56 in endometria of normal controls (n = 6) and IUA patients (n = 6). Scale bars: 100 μm. (H) The correlation analysis of the proportion of eNK cells and ACTA2+ fibroblasts. Error bars, mean ± SD. \*\*\**P* < 0.001, \*\*\*\**P* < 0.0001.

We performed KEGG analysis of differentially expressed genes (DEGs) of eNK cells between normal controls and IUA patients (fold change ≥ 1.5, P < 0.05), and found DEGs were enriched in “Natural killer cell mediated cytoxicity,” “p53 signaling pathway,” “Apoptosis”, which might explain the reduction in the eNK (Supplementary Fig. 1F).

### Characteristics and heterogeneity of eNK cells in the proliferative endometria

We further captured eNK cells and subclustered them into 3 clusters according to the most upregulated genes (Fig. 2A, Supplementary Fig. 2A, B). In normal endometria, cluster 1 was the most abundant eNK subsets, accounting for 42.51% of eNK cells, followed by cluster 2 (31.53%) and cluster 3 (25.96%) (Fig. 2B, Supplementary Fig. 2C).

**Fig. 2.**
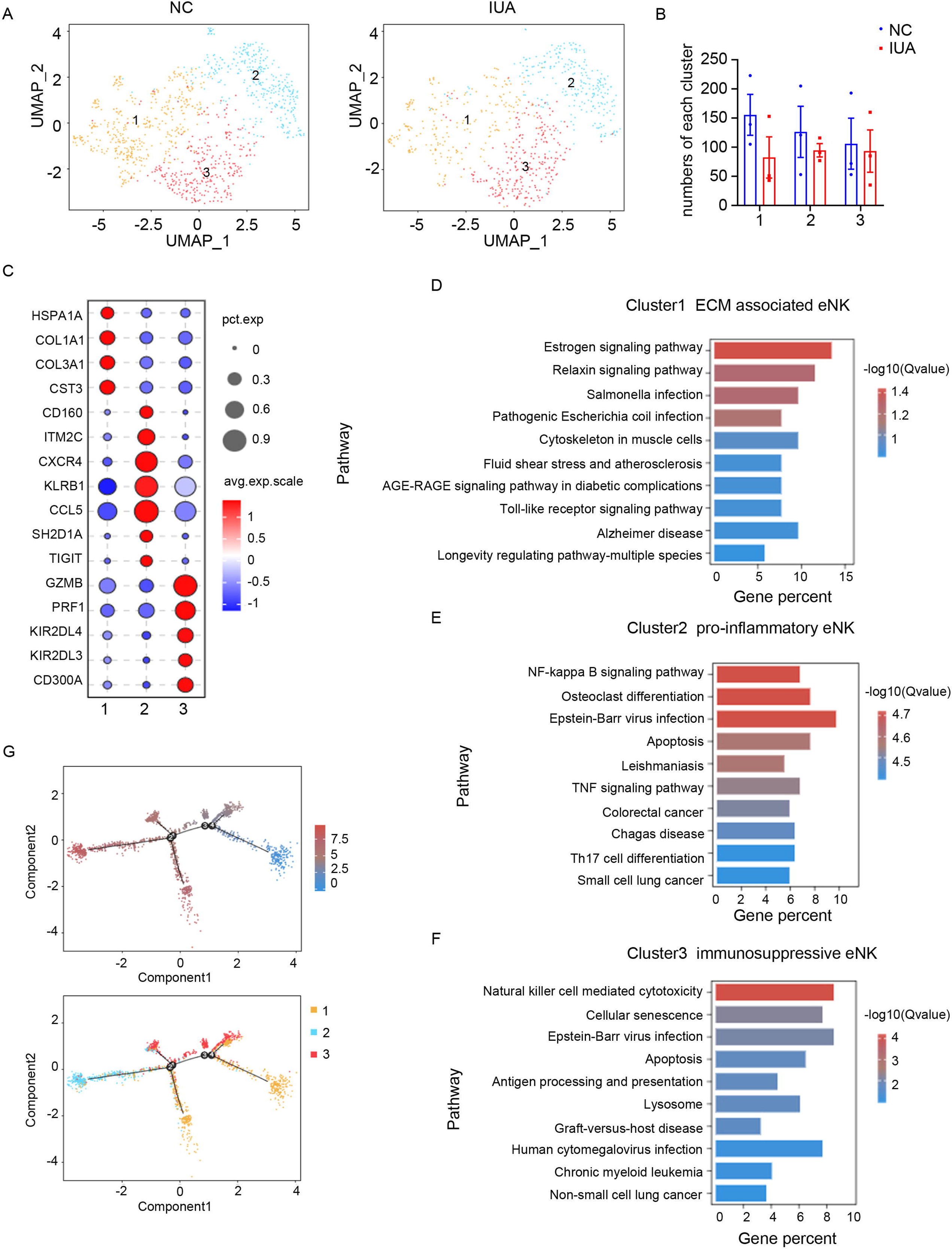
Characteristics and heterogeneity of eNK cells in the proliferating endometria. (A) The UMAP plot showing the distribution of subsets of eNK cells in normal and IUA endometria. (B) The number of each cluster of eNK cells in normal and IUA patients. (C) Bubble diagram showing the expression of marker genes of each cell type. (D-F) The KEGG analysis of the up-regulated genes in three eNK subclusters. (G) Pseudotime analysis of the subsets of eNK cells inferred by Monocle2.

Cluster 1 eNK cells highly expressed ECM components (including COL1A1, COL3A1, ECM1), HSPA1A and CST3 (Fig. 2C). And compared with other clusters, cluster 1 had fewer highly expressed DEGs and lacked specific surface markers (Supplementary Fig. 2A). Hence, we named cluster 1 as “ECM associated eNK”. The KEGG analysis of the DEGs in cluster 1 were enriched in “Estrogen signaling pathway,” “Relaxin signaling pathway,” “Cytoskeleton in muscle cells” (Fig. 2D). Cluster 2 eNK cells were defined by high expression of CXCR4, CD160, KLRB1 and CCL5 (Fig. 2C), and the DEGs in cluster 2 were enriched in “NF-kappa B signaling pathway,” “TNF signaling pathway,” suggesting that cluster 2 could be involved in response to chemokines and promote inflammation (Fig. 2E). We therefore classified cluster 2 as “pro-inflammatory eNK”. We also identified a subset of “immunosuppressive eNK” (cluster 3), marked by high levels of inhibitory receptors, including KIRs, CD300A and granule proteins, including perforin 1 (PRF1), GZMB, granulysin (GNLY) and NKG7 (Fig. 2C). The KEGG analysis of the DEGs in cluster 3 were enriched in “Natural killer cell mediated cytotoxicity” (Fig. 2F).

Pseudotime analysis of eNK cells revealed that “ECM associated eNK” was at initial stage and further differentiated to “pro-inflammatory eNK” and “immunosuppressive eNK” (Fig. 2G). In IUA endometria, all three subsets were decreased inordinately and cluster 1 reduced most significantly (Fig. 2A, B).

Compared with pbNK cells, the receptor expression pattern of eNK cells was closer to that of dNK cells. As shown in Supplementary Fig. 2D, these cells expressed low levels of FCGR3A, ITGA2, and rarely expressed B3GAT1 and KLRG1, which were highly expressed in activated pbNK cells. On the contrary, they expressed relative more CD9 and ITGA1, which were expressed in tissue-resident NK cells, and high levels of inhibitory receptors (KLRC1, KLRD1). However, they were also different from dNK cells in that they expressed fewer NCRs, KIRs and KLRK1. Regarding to the effector molecules, eNK cells expressed abundant genes associated with cytotoxicity, such as NKG7, GNLY, PRF1, granzymes, but lacked the expression of IFNG and TNF.

### eNK cells maintain ESCs homeostasis and promote dedifferentiation of myofibroblasts

Due to the lack of surface marker molecules in “ECM associated eNK” subgroup, it is difficult to isolate this group of cells individually. Since cluster 1 subgroup was the largest population and showed the most significant reduction, and was the progenitor cells of other two subgroups of eNK cells, we isolated total eNK cells to investigate their effect on endometrial fibrosis. eNK cells were sorted by CD56 magnetic beads after digesting and isolating normal human endometrial cells, and the purity was more than 95% identified by the flow cytometry (Supplementary Fig. 3A, B). As shown in Supplementary Fig. 3C-E, when ESCs proliferated and passaged into the third passage in vitro, they exhibited morphological changes from spindle to stellate and the expression of α-SMA and Collagen 1 increased, which suggested the differentiation from ESCs to myofibroblasts during the proliferation of ESCs. However, if co-cultured ESCs with eNK cells, the fibrotic markers, α-SMA, Collagen 1 and Fibronectin were significantly reduced at the third passage of ESCs, indicating that eNK cells inhibited ESCs differentiation into myofibroblasts during their proliferation to maintain endometrial homeostasis (Fig. 3A-E).

**Fig. 3.**
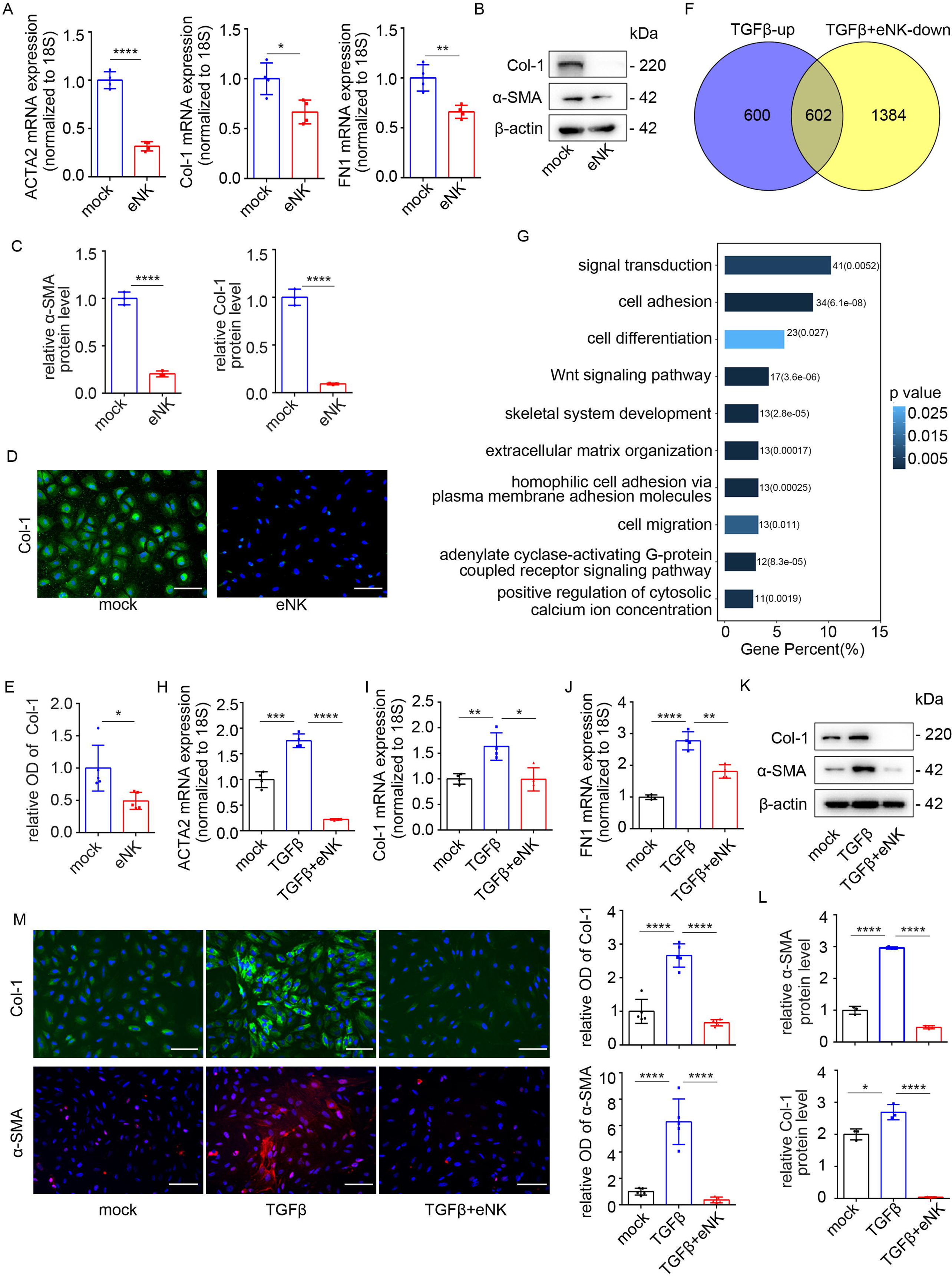
eNK cells can maintain ESCs homeostasis and promote the dedifferentiation of TGFβ1-induced myofibroblasts. (A) The mRNA levels of ACTA2 (α-SMA), Col-1 (Collagen 1) and FN1 (Fibronectin 1) in ESCs (n = 4) treated with the supernatants of eNK cells for 48 h detected by qRT-PCR. (B) The protein levels of α-SMA and Col-1 in ESCs (n = 3) treated with supernatants of eNK cells for 48 h detected by Western Blotting. (C) Relative band intensities of α-SMA and Col-1 analyzed by Image J. (D) Immunofluorescence staining of Col-1 in ESCs (n = 5) treated with the supernatants of eNK cells for 48 h. Scale bars: 100 μm. (E) The relative optical densities (OD) of Col-1 staining analyzed by Image J. (F-L) The ESCs were pr-treated with 10 ng/ml TGFβ1 for 48 h and then incubated with supernatants of eNK cells. (F) Venn analysis of the up-regulated genes in TGFβ1-induced myofibroblasts and down-regulated genes in eNK-treated myofibroblasts. (G) GO biological process enrichment analysis of the screened genes. (H-J) The mRNA levels of ACTA2, Col-1 and FN1 in ESCs (n = 4) detected by qRT-PCR. (K) The protein levels of α-SMA and Col-1 in ESCs (n = 3) detected by Western Blotting. (L) Relative band intensities of α-SMA and Col-1 analyzed by Image J. (M) Immunofluorescence staining of α-SMA and Col-1 in ESCs (n = 5) and their relative optical densities (OD) were analyzed by Image J. Scale bars: 100 μm. Error bars, mean ± SD. \**P* < 0.05, \*\**P* < 0.01, \*\*\**P* < 0.001, \*\*\*\**P* < 0.0001.

We then used TGFβ1 to induce the differentiation of ESCs into myofibroblasts to further investigate the effect of eNK cells on myofibroblasts. The results showed that the morphology of myofibroblasts changed from flat and stellate to spindle shape when co-cultured with eNK cells and we also found that eNK cells exhibited low cytotoxicity to myofibroblasts, despite their high expression of effector molecules including PRF1, GZMB, GNLY (Supplementary Fig. 3F-H). We then performed RNA sequencing (RNA-seq) analysis of ESCs, myofibroblasts and the myofibroblasts co-cultured with eNK cells. Venn analysis of the upregulated genes in myofibroblasts and downregulated genes in eNK-treated myofibroblasts showed that 602 genes were filtered out (Fig. 3F). GO biological process enrichment analysis of these 602 differentially genes revealed that fibrosis-related terms, including “cell adhesion,” “Wnt signaling pathway,” “skeletal system development,” “extracellular matrix organization” and “cell migration”, were upregulated in myofibroblasts and downregulated in eNK-treated myofibroblasts (Fig. 3G). To verify the result of RNA-seq, we examined the mRNA and protein levels of fibrotic markers, including α-SMA, Collagen 1 and Fibronectin, which were significantly down-regulated in eNK co-cultured myofibroblasts, indicating that eNK cells could promote the dedifferentiation of myofibroblasts (Fig. 3H-M).

### PGD2 is an important effector of eNK cells in maintaining homeostasis of ESCs and promoting myofibroblasts dedifferentiation

To understand the molecular mechanism by which eNK cells maintained endometrial stromal compartment homeostasis and promoted myofibroblasts dedifferentiation, we further compared DEGs of eNK cells between IUA endometria and normal controls. The results showed that the prostaglandin D2 synthetase (PTGDS) was significantly decreased in IUA patients, which was confirmed by immunofluorescence staining of CD56 and PTGDS in endometria (Fig. 4A, B). Since PTGDS is a catalytic enzyme to synthesize PGD2 [23], and PGD2 was detected in the supernatant of eNK cells isolated from proliferative endometria (Fig. 4C), we then explored the role of PGD2 in endometrial homeostasis and endometrial fibrosis. The results showed that in the presence of PGD2, the fibrotic markers (α-SMA, Collagen 1 and Fibronectin) of ESCs were significantly down-regulated during in vitro passage (Supplementary Fig. 4A-C). Further, we used PGD2 to treat TGFβ1-induced myofibroblasts and found that α-SMA, Collagen 1 and Fibronectin were also significantly decreased in both mRNA and protein levels (Fig. 4D-H). However, pbNK cells produced almost no PGD2 (Fig. 4C) and had no role in inhibiting the differentiation of ESCs into myofibroblasts and nor in promoting the dedifferentiation of myofibroblasts (Supplementary Fig. 4D, E). To further determine whether eNK cells inhibit endometrial fibrosis by secreting PGD2, we next pretreated myofibroblasts with an inhibitor of PGD2 receptor, PGD2-IN-1, and the results showed that the effect of PGD2 or eNK cells on the myofibroblasts dedifferentiation nearly disappeared (Supplementary Fig. 4F, Fig. 4I-M). And to exclude the effect of PGD2-IN-1 itself on myofibroblasts, we treated myofibroblasts only with PGD2-IN-1, and found that no significant changes were observed in the protein levels of α-SMA, Collagen 1 (Supplementary Fig. 4G). Collectively, these results suggested that PGD2 produced by eNK cells could be a critical player in maintaining endometrial homeostasis and inhibiting endometrial fibrosis.

**Fig. 4.**
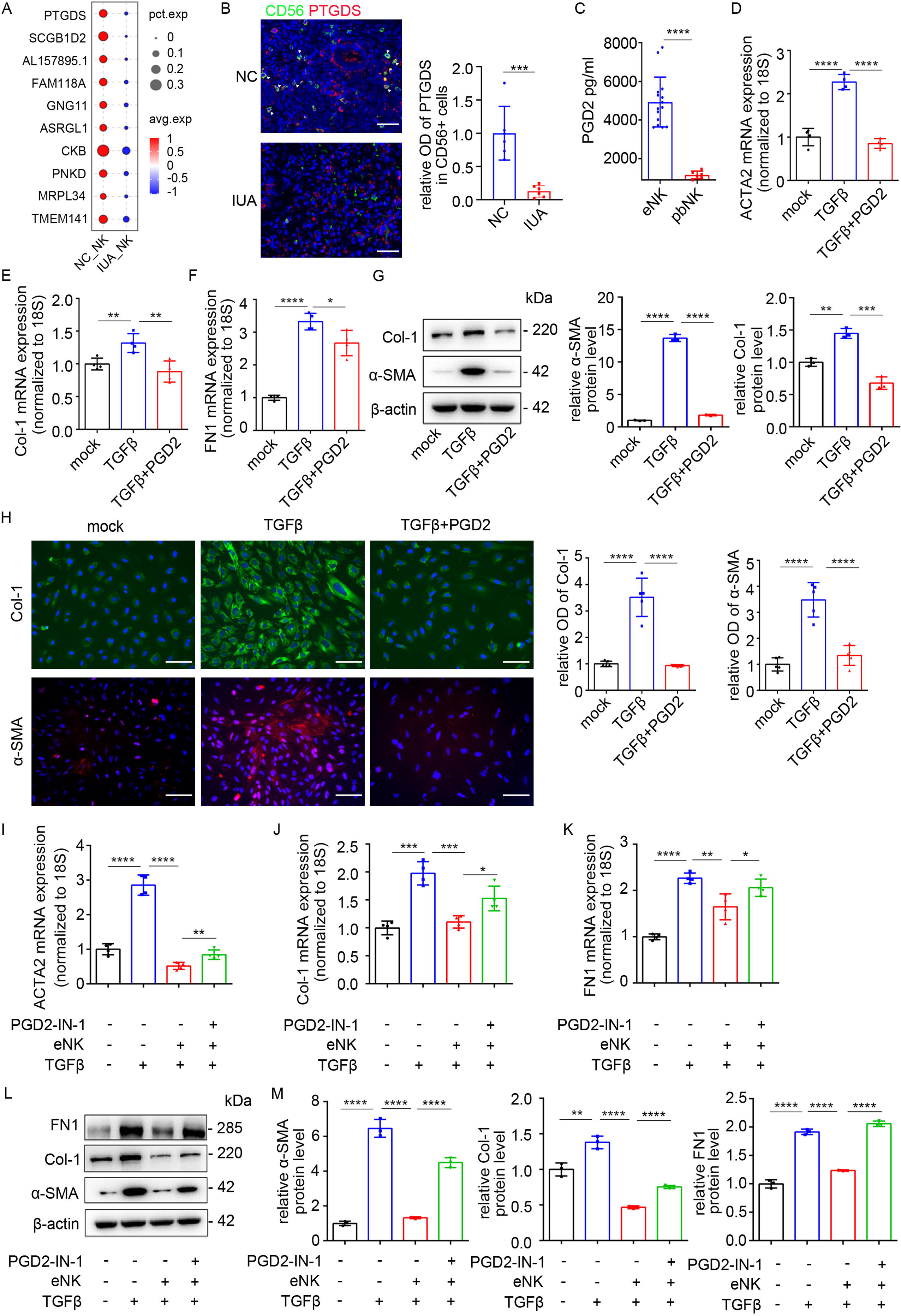
PGD2 mediates the dedifferentiation of eNK cells on TGFβ1-induced myofibroblasts. (A) Bubble diagram showing the top10 DEGs of eNK cells between normal controls and IUA patients based on scRNA-seq analysis. (B) Immunofluorescence staining of CD56 (green) and PTGDS (red) in endometria of normal controls (n = 6) and IUA patients (n = 6) and the relative optical densities (OD) of PTGDS staining in CD56+ cells analyzed by Image J. Scale bars: 50 μm. (C) The level of PGD2 in the supernatants of eNK cells (n = 15) and pbNK cells (n = 6). (D-H) The ESCs were pre-treated with 10 ng/ml TGFβ1 for 48 h and then treated with 10 μM PGD2 for 48 h. (D-F) The mRNA levels of ACTA2, Col-1 and FN1 in ESCs (n = 4) examined by qRT-PCR. (G) Left: The protein levels of α-SMA and Col-1 in ESCs (n = 3) examined by Western Blotting. Right: Relative band intensities analyzed by Image J. (H) Immunofluorescence staining of α-SMA and Col-1 in ESCs (n = 5) and their relative optical densities (OD) were analyzed by Image J. Scale bars: 100 μm. (I-M) The ESCs were treated with 1 μM PGD2-IN-1 24 h before eNK cells incubation. (I-K) The mRNA levels of ACTA2, Col-1 and FN1 in ESCs (n = 4) examined by qRT-PCR. (L) The protein levels of α-SMA, Col-1 and FN1 in ESCs (n = 3) examined by Western Blotting. (M) Relative band intensities of α-SMA, Col-1 and FN1 analyzed by Image J. Error bars, mean ± SD. \**P* < 0.05, \*\**P* < 0.01, \*\*\**P* < 0.001, \*\*\*\**P* < 0.0001.

### ZNF521 is the main target molecule of PGD2 against endometrial fibrosis

To investigate the molecular mechanism of eNK cells and PGD2 in maintaining ESCs homeostasis and promoting myofibroblasts dedifferentiation, we conducted Venn analysis of the 602 genes and myofibroblasts associated transcription factors (TF) [24]. 5 TFs were filtered out, including BNC2, OSR1, ZNF521, FOS and BCL11A (Fig. 5A, B). We validated the changes in the expression levels of these genes in vitro and found that only the mRNA and protein levels of ZNF521 were up-regulated in myofibroblasts and down-regulated after eNK cells or PGD2 treatment among the 5 TFs (Fig. 5C-E, Supplementary Fig. 5A-F). Then we detected the expression of ZNF521 in endometria of IUA patients, and found that ZNF521 was significantly increased in IUA endometria (Fig. 5F, G).

**Fig. 5.**
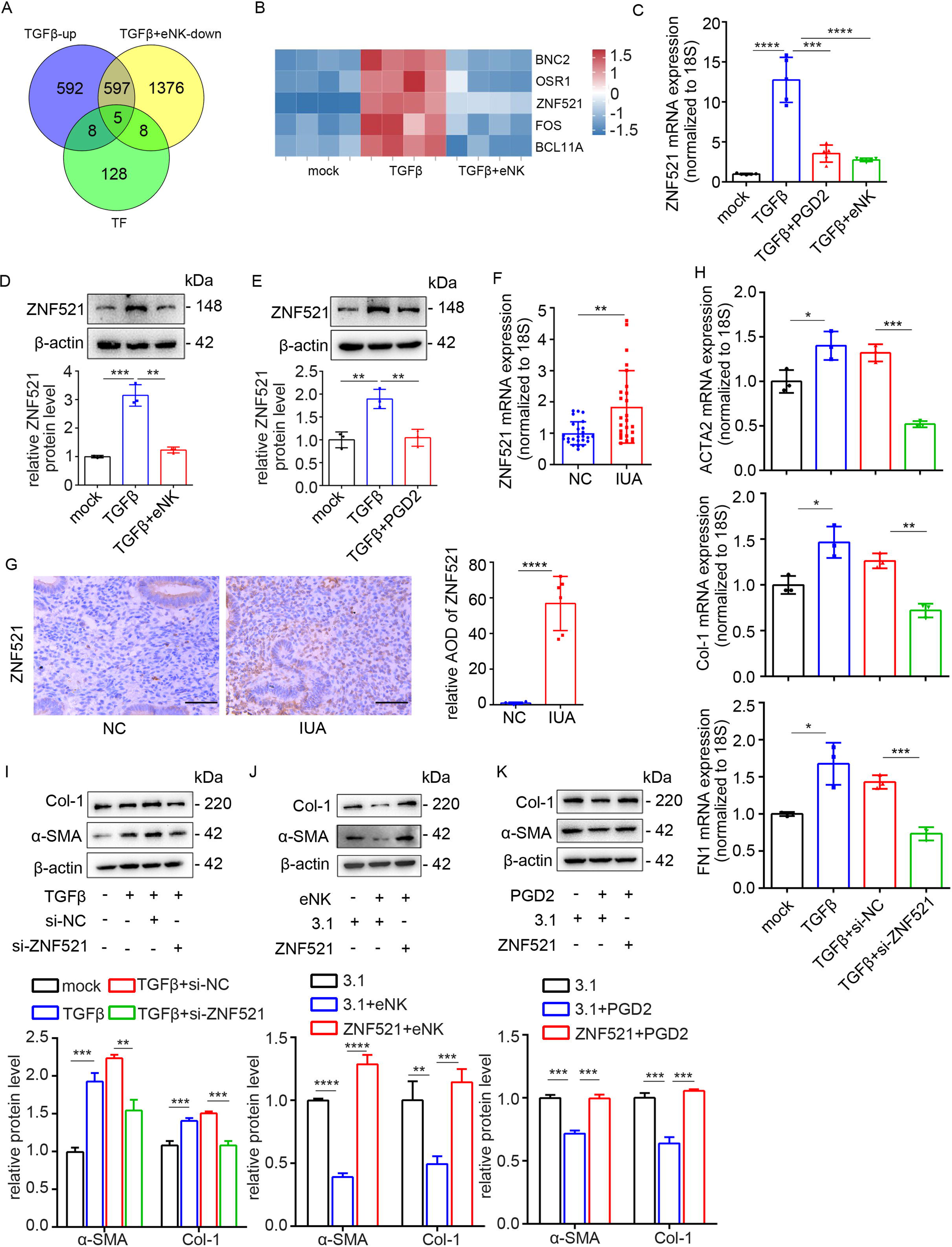
eNK cells or PGD2 promote myofibroblast dedifferentiation via regulating ZNF521. (A) Venn analysis of the genes upregulated in myofibroblasts, downregulated in eNK-treated myofibroblasts and transcription factors associated with myofibroblasts. (B) Heatmap of genes (BNC2, OSR1, ZNF521, FOS and BCL11A) in ESCs, myofibroblasts and eNK-treated myofibroblasts. (C) The mRNA level of ZNF521 in ESCs (n = 5) detected by qRT-PCR. (D-E) Top: The protein level of ZNF521 in ESCs (n = 3). Bottom: Relative band intensities analyzed by Image J. (F) The mRNA level of ZNF521 in endometrial of normal controls (n = 25) and IUA patients (n = 25) detected by qRT-PCR. (G) Immunohistochemical staining of ZNF521 in endometria of normal controls (n = 6) and IUA patients (n = 6). Scale bars: 50 μm. (H) The mRNA levels of ACTA2, Col-1 and FN1 in ESCs (n = 3, pre-treated with TGFβ1) transfected with si-NC or si-ZNF521 for 48 h examined by qRT-PCR. (I) Top: The protein levels of α-SMA and Col-1 in ESCs (n = 3) detected by Western Blotting. Bottom: Relative band intensities analyzed by Image J. (J) Top: The protein levels of α-SMA and Col-1 in ESCs (n = 3, pre-treated with eNK cells) transfected with ZNF521 plasmid for 48 h detected by Western Blotting. Bottom: Relative band intensities analyzed by Image J. (K) Top: The protein levels of α-SMA and Col-1 in ESCs (n = 3, pre-treated with PGD2) transfected with ZNF521 plasmid detected by Western Blotting. Bottom: Relative band intensities analyzed by Image J. Error bars, mean ± SD. \**P* < 0.05, \*\**P* < 0.01, \*\*\**P* < 0.001, \*\*\*\**P* < 0.0001.

ZNF521 belongs to the C2H2 zinc finger protein family with critical roles in hematopoietic progenitor cell homeostasis, neuronal differentiation, bone formation and cancer progression [25–29]. However, its role in endometria and fibrotic diseases remains unclear. To investigate the role of ZNF521 in endometrial fibrosis, we performed the suppression and overexpression of ZNF521 in ESCs. The results showed that the mRNA and protein expression of ZNF521 were significantly decreased after silencing ZNF521 by siRNA transfection, along with the downregulation of α-SMA, Collagen 1 and Fibronectin (Supplementary Fig. 5G-J). We transfected ESCs with a plasmid containing ZNF521 cDNA and confirmed that the mRNA and protein expression of ZNF521 were increased (Supplementary Fig. 5K, L), and meanwhile the expression of α-SMA, Collagen 1 and Fibronectin were increased compared to control plasmids (Supplementary Fig. 5M, N). Furthermore, we transfected siZNF521 into ESCs pre-stimulated with TGFβ1 and the fibrotic markers were reduced (Fig. 5H, I). Importantly, the cytoprotective effect of eNK cells or PGD2 was lost in the ZNF521-overexpression ESCs, suggesting that the function of eNK cells or PGD2 in maintaining ESCs homeostasis and promoting myofibroblasts dedifferentiation was through regulating ZNF521 (Fig. 5J, K).

### eNK cells ameliorate endometrial fibrosis in IUA murine model through PGD2 down-modulation of ZNF521

To confirm the role of eNK cells against endometrial fibrosis in vivo, we generated a mechanical injury-induced IUA-like mice model by curettage as previously reported [30]. Compared with the sham-operated group, the IUA group had more collagen deposition and myofibroblasts as indicated by Masson staining and IHC staining of α-SMA and Collagen I (Supplementary Fig. 6A-D). In addition, the expression of ZNF521 was significantly increased in endometria of IUA-like mice (Supplementary Fig. 6E).

Then we applied this IUA-like mice model to assess the anti-fibrotic effects of eNK cells and PGD2. We injected conditioned medium of eNK cells or PGD2 or blank medium intraperitoneally once a day for 10 days after the second curettage and then fibrosis-related markers were evaluated (Fig. 6A). Compared with blank medium administration group, the collagen deposition in eNK cells-treated group was significantly alleviated, and the expression level of α-SMA, Collagen I and ZNF521 were also downregulated. And similar results were found in PGD2-treated group (Fig. 6B, C). To further confirm the role of PGD2 in conditioned medium of eNK cells, we administrated PGD2-IN-1 together. Masson staining and immunohistochemical staining of α-SMA and Collagen I showed that the protective effect of eNK cells in the PGD2-IN-1-treated group largely disappeared compared with eNK-treated group (Fig. 6B, C). Furthermore, in conditioned medium of pbNK cells-treated group, which barely secreted PGD2, we couldn’t find any attenuation of endometrial fibrosis compared with IUA group (Fig. 6B, C).

**Fig. 6.**
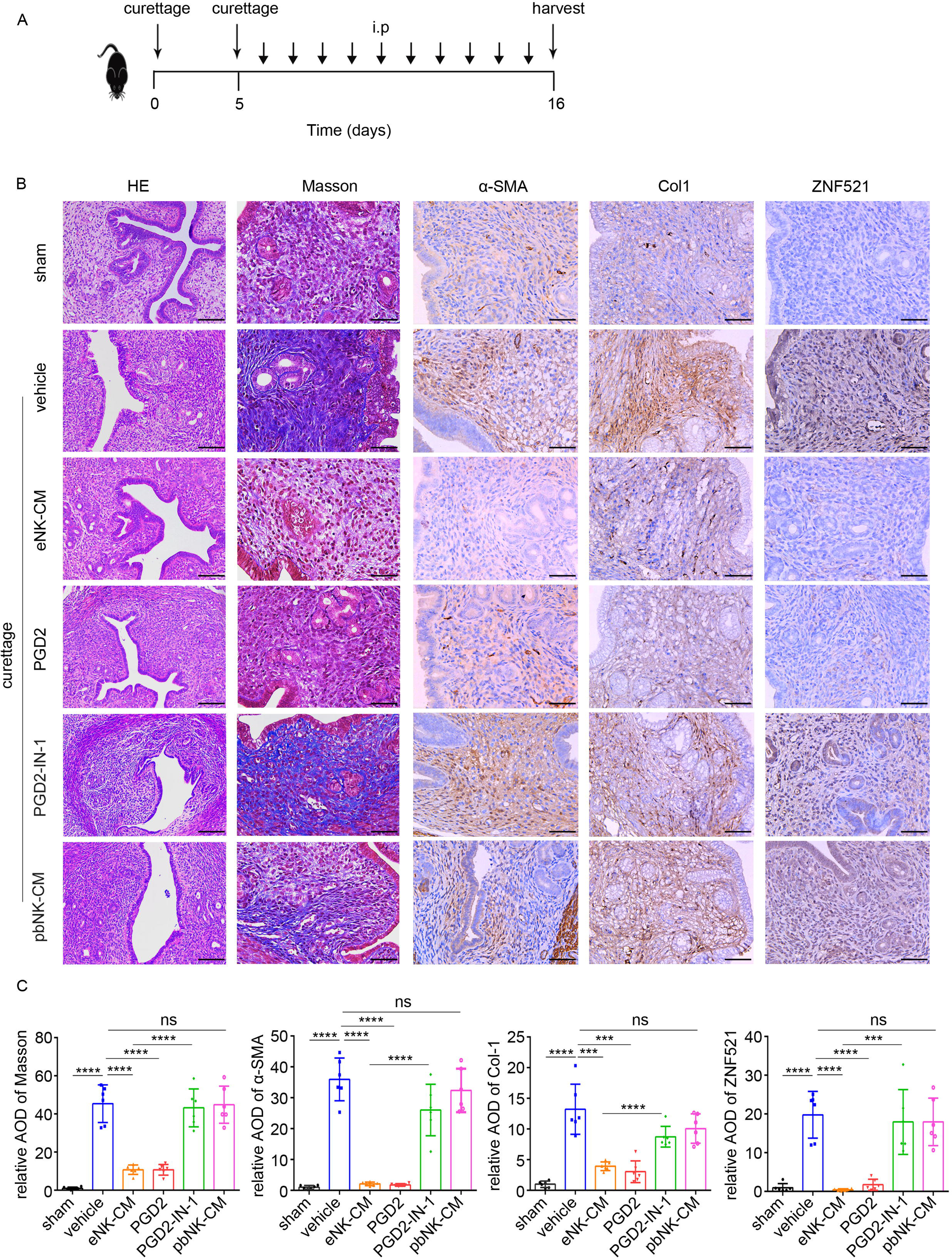
eNK cells or PGD2 attenuate endometrial fibrosis in vivo. (A) The mice were divided into the following groups: Sham operation (n = 6), curettage (n = 6), curettage + supernatants of eNK cells (n = 6), curettage + PGD2 (200 ug/kg, n = 6), curettage + supernatants of eNK cells + PGD2-IN-1 (250 ug/kg, n = 6), curettage + supernatants of pbNK cells (n = 6). (B) HE (scale bars of HE: 100 μm), Masson staining and immunohistochemical staining of α-SMA, Col-1 and ZNF521 in endometria of mice. Scale bars: 50 μm. (C) The relative AOD of Masson, α-SMA, Col-1 and ZNF521 analyzed by Image J. Error bars, mean ± SD. \*\*\**P* < 0.001, \*\*\*\**P* < 0.0001, ns: not significant.

## Discussion

The human endometrium is a special magical tissue consisted of a variety of non-immune cells and immune cells [22, 31]. Different from endometrial fibrosis in IUA patients, the endometria in normal women of childbearing age undergo about 400 cycles of proliferation, differentiation, shedding and repair without fibrosis [32]. However, the mechanism of maintaining endometrial cell homeostasis remains unclear. It has been speculated that this may be the result of the delicate interplay among various cellular components, especially the regulatory role played by immune cells in the endometrium [31–33]. IUA is a common and refractory uterine infertility characterized by endometrial fibrosis to which the differentiation of the largest proportion of ESCs into myofibroblasts contributes [34]. In this study, we demonstrated a significant decrease in eNK cells in late proliferative endometria of IUA patients, accompanied by an increase in myofibroblasts. There was a highly negative correlation between eNK cells and myofibroblasts (Fig. 1H). We further showed that eNK cells could inhibit the differentiation of ESCs into myofibroblasts and promote the latter dedifferentiation to prevent the endometria from fibrosis, in which eNK cells acted mainly by downregulating ZNF521 through secretion of PGD2.

It is well-described that not only adaptive lymphocytes undergo differentiation in response to environmental cues but also innate immune cells including uterine NK cells are subject to regulated differentiation [17]. In uterine, NK cells reside in the endometrium where non-immune cells experience periodic proliferation and differentiation under the regulation of estrogen and progesterone [16, 17, 35]. In response to progesterone after ovulation (i.e. secretory phase), ESCs differentiate into decidual stromal cells to secrete a large amount of interleukin-15 (IL-15), which promotes the rapid proliferation and recruitment of NK cells [16, 17]. Accumulating evidence has demonstrated the powerful immunomodulatory function of dNK cells to achieve immune tolerance at the maternal-fetal interface and besides, dNK cells can promote fetal and placental development [18–20, 36, 37]. But so far, it is not clear what is the function and phenotype of eNK cells in the proliferative phase before progesterone arrives. In this study, we demonstrated that eNK cells exhibited three distinct subtypes and they were different from pbNK cells and dNK cells in both phenotypes and functionalities (Supplementary Fig 2D). Among them, cluster 1 eNK cells, not being reported before, accounted for the largest proportion and mainly expressed ECM-related molecules. The DEGs in cluster 2 eNK cells were enriched in inflammation-related pathways, implicating a pro-inflammatory role. And although cluster 3 cells highly expressed a variety of granzymes and perforin, they highly expressed multiple inhibitory receptors at the same time, which might explain the low cytotoxicity to target cells. Our study indicated that eNK cells played an important role in inhibiting differentiation of ESCs into myofibroblasts and promoting myofibroblasts dedifferentiation to prevent endometrial fibrosis, although we were not able to observe their function of the different subgroups separately.

Tissue fibrosis is the most common cause of loss of normal tissue functions. In the process of exploring the therapeutic strategies for fibrotic diseases, NK cells have attracted more and more attention as a new means [13, 38–40]. In liver fibrosis, NK cells can induce the apoptosis of activated hepatic stellate cells through cytotoxicity and IFNγ secretion to alleviate fibrosis [13, 38, 39]. In autoimmune myocarditis, NK cells can limit eosinophil accumulation to inhibit cardiac fibrosis [40]. Here we demonstrated that eNK cells exhibited an anti-fibrotic function in endometria by a distinct and intriguing way and they presented low cytotoxicity towards myofibroblasts and produced only minimal amounts of IFN-γ. Instead, eNK cells inhibited endometrial fibrosis by restraining ESCs differentiation into myofibroblasts and promoting the dedifferentiation of myofibroblasts through secreting PGD2. This unique mechanism underscores that the anti-fibrotic effect of eNK cells has tissue specificity.

PGD2 is a bioactive lipid mediator specifically catalyzed by PTGDS and the expression level of PTGDS determines the level of PGD2 [41]. PGD2 secreted by eNK cells was detected in the culture medium of normal eNK cells (Fig. 4C). Due to few eNK cells in IUA endometria, it was difficult to isolate eNK cells to detect the secretion of PGD2, so we used immunofluorescence staining and found that the expression of PTGDS in eNK cells was significantly decreased in IUA endometria (Fig. 4B). Similar to eNK cells, PGD2 could also inhibit the differentiation of ESCs into myofibroblasts and promote the dedifferentiation of myofibroblasts to inhibit endometrial fibrosis. Furthermore, we delved into the molecular mechanisms underlying the inhibitory effect of eNK cells-derived PGD2 on endometrial fibrosis and identified a novel molecule, ZNF521. Although ZNF521 has been linked to tumor progression and epithelial-mesenchymal transition [27, 42], its role in fibrosis is unclear. Our study unveiled the ZNF521’s pro-fibrotic effect in endometria and eNK cells-derived PGD2 inhibited ZNF521 expression to perform the function of anti-endometrial fibrosis.

This study has some limitations. Firstly, the lack of specific surface markers for “ECM associated eNK” hinders our ability to isolate this subset for further investigations. More direct evidence is needed to elucidate the functions of “ECM associated eNK” and other eNK subsets in future. Additionally, given the differences in cell characteristics and the ongoing debate regarding the origin and phenotype of NK cells in the uterus of non-pregnant mice [43, 44], we were unable to establish an animal model of eNK cells deficiency. However, by constructing an IUA-like mouse model and treating it with supernatants from eNK cells or PGD2 or an inhibitor, we confirmed that eNK cells and their secreted PGD2 played crucial roles in preventing endometrial fibrosis.

In conclusion and clinical implication, our data highlight the critical role of eNK cells in maintaining endometrial homeostasis and preventing endometria from fibrosis, which shed light to IUA therapy.

## Materials and Methods

### Human samples

The human endometrial tissues and peripheral blood samples involved in this study were approved by the Ethics Committee of Nanjing Drum Tower Hospital, Affiliated Hospital of Nanjing University Medical School (SC2019-002-02) and written informed consent was obtained from all participants. The endometrial biopsies were obtained from child-bearing women undergoing hysteroscopy at Nanjing Drum Tower Hospital and all patients involved were in the late proliferative phase of the menstrual cycle as previously reported [6, 30]. Fresh endometrial samples were obtained from patients with severe IUA (n=25) as classified by the American Fertility Society [45] or non-IUA patients (n=25) who had tubal infertility with normal ovary function, normal menstrual blood volume, and endometrium thickness. Women with a history of tuberculosis, vaginitis, chronic endometritis and hormone therapy were excluded. Peripheral blood samples were collected from 6 healthy donors. The clinical characteristics of normal controls and IUA patients were summarized in Table 1.

**Table 1.** Clinical characteristics of normal controls and IUA patients.

| Parameter | Control (n=25) | IUA (n=25) | P value |
| --- | --- | --- | --- |
| Age (y) | 32.12 ± 3.91 | 30.40 ± 3.54 | >0.05 |
| IUA score (AFS) | 0 | 9.16 ± 1.34 | <0.05 |
| Endometrial thickness (mm) | 10.62 ± 3.04 | 5.24 ± 1.52 | <0.05 |

AFS: American Fertility Society, 1988.

### Isolation of primary endometrial stromal cells and NK cells

The fresh human endometrial tissue samples were rinsed with PBS to remove any visible blood clots. Subsequently, they were minced into smaller pieces and digested with 0.1% trypsin (Wisent Inc.) for 10 min and then transferred to a solution of 0.8 mg/ml collagenase type I (Sigma) for 90 min with gentle shaking in a humidified incubator at 37L and 5% CO2. ESCs were obtained through a 40 μm cell strainer (BD Falcon) to separate from epithelial glands. ESCs were cultured in DMEM/F12 (Wisent Inc) supplemented with 10% FBS (Gibco) and 1% penicillin-streptomycin (Wisent Inc). For eNK cells isolation, mononuclear cells extracted from endometrial samples after digestion were suspended with 30% Percoll (Cytiva) and then added to liquid surface of 70% Percoll carefully, centrifuged at 900 g for 30 min to remove cell debris. The eNK cells were further purified by EasySep™ Human CD56 Positive Selection Kit II (STEMCELL). Only eNK cells (FVS510-CD56+CD3-) of purity > 95% were employed for each assay. For pbNK cells isolation, peripheral blood mononuclear cells were obtained by density gradient centrifugation through Ficoll (Cytiva). Then NK cells were further purified by magnetic bead sorting. eNK cells and pbNK cells were cultured in OptiVitro NK cell expansion kit (Excell Bio).

### RNA isolation and quantitative real-time PCR (qRT-PCR)

Total RNA from cells or frozen tissues was extracted using TRNzol reagent (Tiangen) and trichloromethane, then an equal volume of isopropanol was added to precipitate RNA. The quality and purity of RNA was assessed by Nano Drop spectroscopy (Thermo Fisher Scientific). cDNA was reverse transcribed using a HiScript III 1st Strand cDNA Synthesis Kit (Vazyme) and qPCR was performed using ChamQ SYBR® qPCR Master Mix (Vazyme) with a LightCycler 96 instrument (Roche). The relative mRNA expression was calculated by the value of 2^-ΔCt^ normalized to 18S. Gene primers used in this study are listed in Table 2.

**Table 2.**
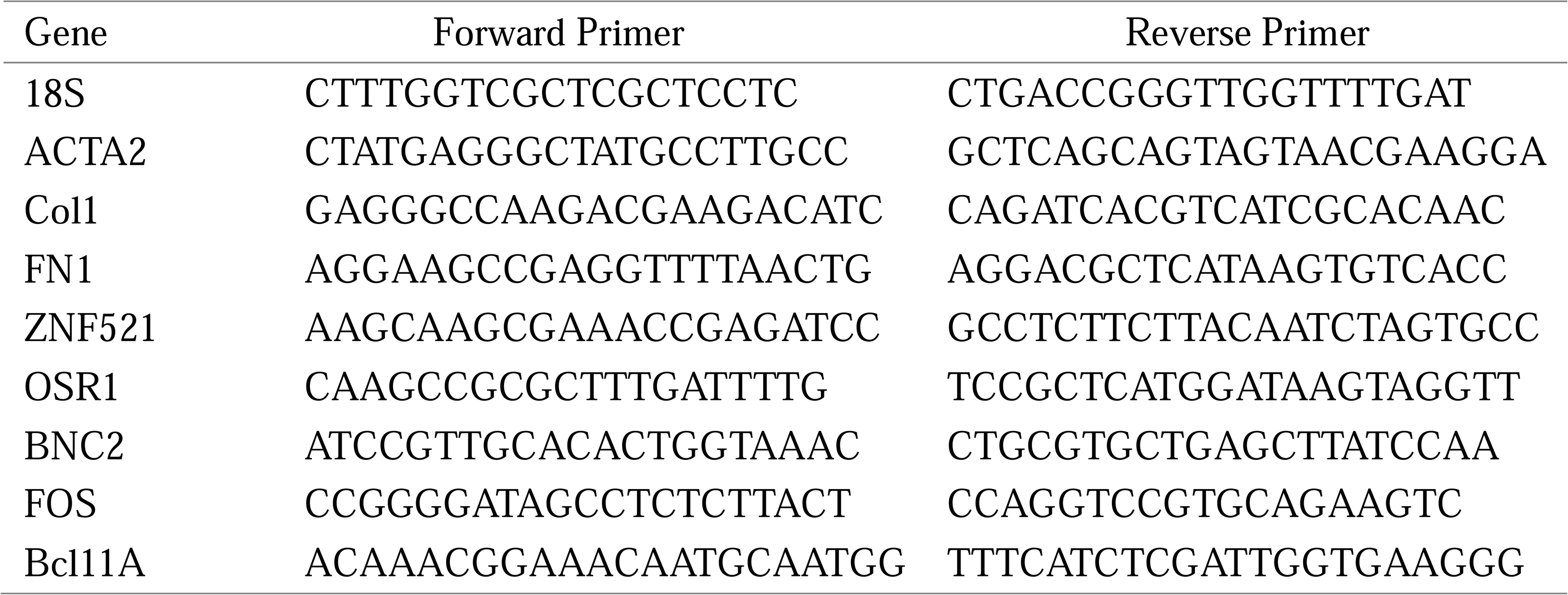
Primer sequences used in this study.

### Flow cytometry assay

For the human samples, single-cell suspensions were prepared as described above and then centrifuged at 300 g for 5 min. The cells were subsequently washed and resuspended with 100 μl PBS. To stain the cells, they were incubated with antibodies, including FVS510 (BD Biosciences), CD45 BV421 (Biolegend), CD3 APC-cy7 (BD Biosciences) and CD56-PE (Biolegend), at room temperature for 20 min. The gating strategy employed to identify human eNK cells was based on the expression pattern FVS510-CD45+CD3-CD56+. Flow cytometry analysis was performed using CytoFLEX instruments (Beckman Coulter) and the acquired data were analyzed with CytExpert v2.3 software (Beckman Coulter). The antibodies used are listed in Table 3.

**Table 3.**
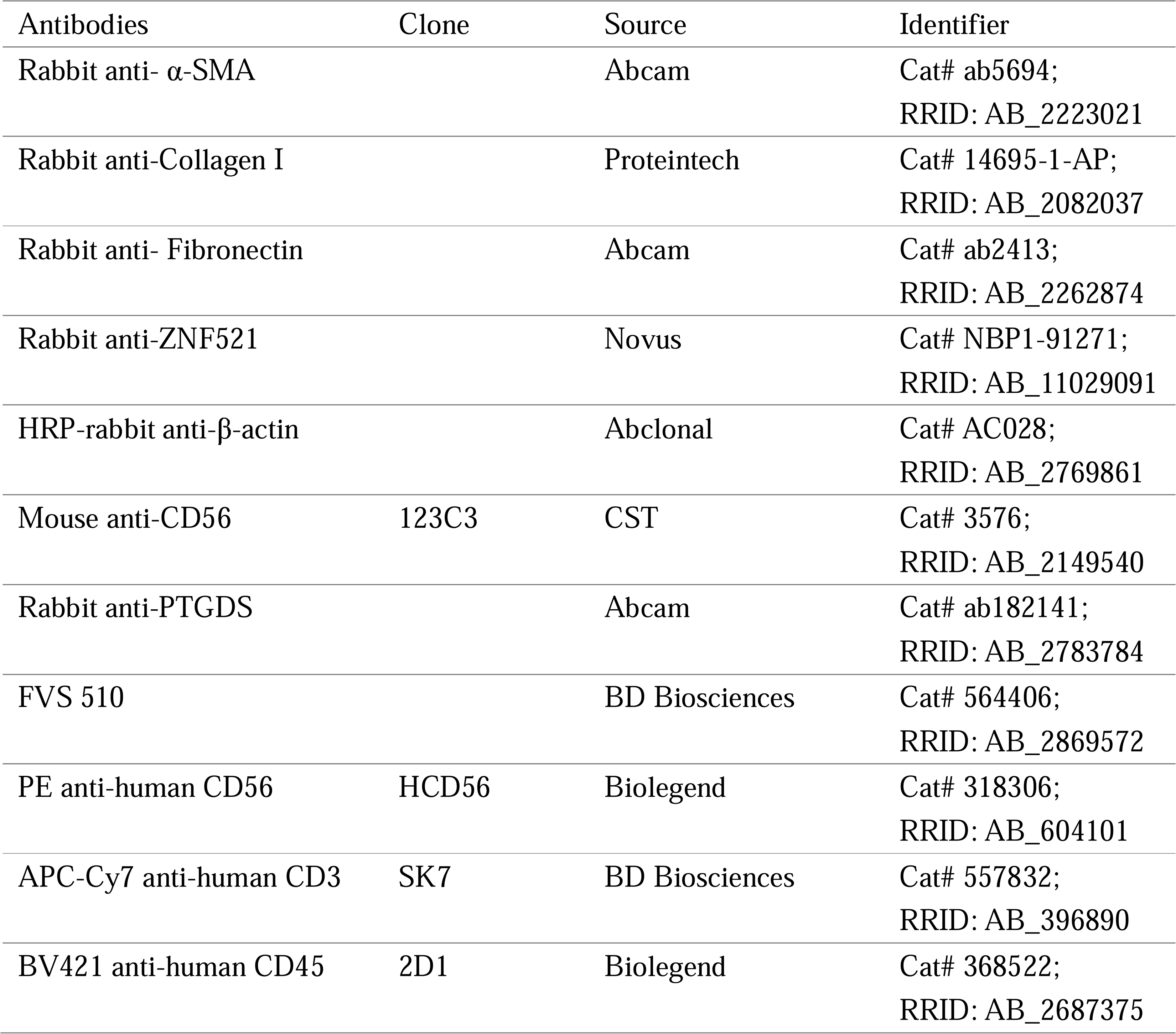
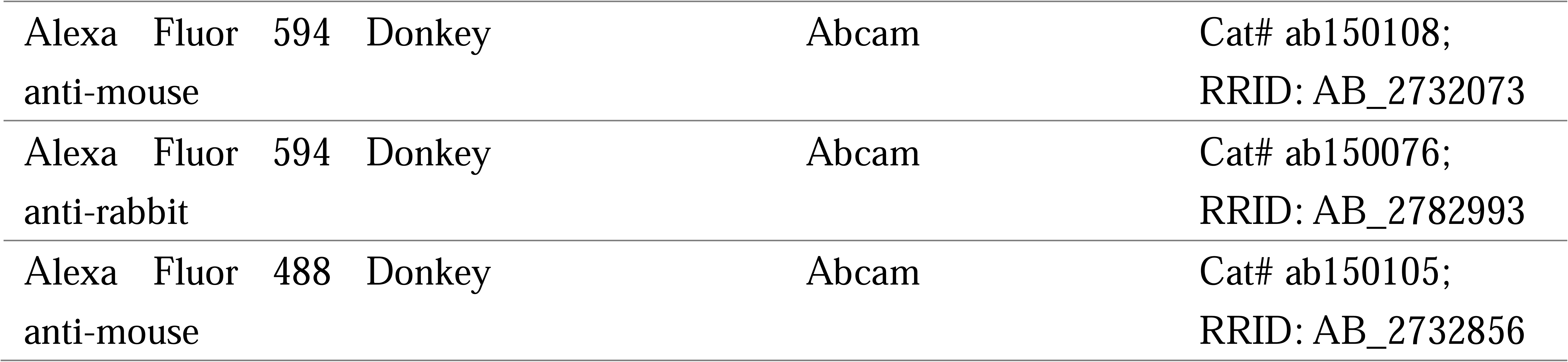
Antibodies used in this study.

### Immunofluorescence

The fresh human endometrial tissues were fixed overnight in 4% paraformaldehyde (PFA), followed by dehydration in a gradient series of sucrose (7.5%, 15% and 30% respectively). The tissues were then embedded in OCT and frozen in liquid nitrogen. OCT-embedded samples were cut into 5 μm thickness. The sections were blocked with 2% BSA for 30 min, incubated with the primary antibodies at 4L overnight, and then with the secondary antibodies away from light for 1 h. Nuclei were stained with DAPI (Abcam). The ESCs were washed with PBS for three times, fixed with 4% PFA for 15 min, permeabilized with 0.1% PBST, blocked with 2% BSA and then the cells were incubated with primary and secondary antibodies, followed by DAPI staining. Images were captured by a fluorescence microscope (DMi8, Leica). The antibodies used are listed in Table 3.

### Histology and immunohistochemistry

The human and mouse tissue samples were fixed with 4% PFA overnight, subsequently dehydrated using a gradient series of alcohol, and embedded in paraffin. The overall morphology was assessed by hematoxylin and eosin staining. To assess collagen deposition, Masson’s Trichrome staining (Solarbio) was performed according to the manufacturer’s instructions. For immunohistochemical staining, the sections were deparaffinized, rehydrated and the endogenous peroxidase was eliminated followed by heat-mediated antigen retrieval. Then the primary and secondary antibodies were added sequentially. The sections were finally developed with DAB substrate to visualize the antigen signals and counterstained with hematoxylin. Images were captured by a microscope (DMi8, Leica). The antibodies used are listed in Table 3.

### Cytotoxicity assay

The cytotoxicity of eNK cells was tested using a lactate dehydrogenase (LDH) release assay according to kit instructions (Beyotime). Briefly, ESCs and TGFβ1-induced myofibroblasts (1×10^4 cells per well) seeding into 96-well plate were employed as targets. Then eNK cells (1×10^5 cells per well) were employed as effectors seeding into the cell culture plate for killing for 24 h before cytotoxicity assay.

### Bulk RNA-sequencing and data analysis

Analysis of differential gene expression profiles in ESCs, myofibroblasts induced by TGFβ1 and myofibroblasts co-cultured with eNK cells was performed on an Illumina NovaSeq6000 by Gene Denovo Biotechnology Co. (Guangzhou). Total RNA was extracted by TRIzol reagent. Differential expression analysis between the various groups was performed using the DESeq2 software. Genes were deemed as differentially expressed genes (DEGs) if they met the criteria of having a Q value < 0.05 and absolute fold change > 2. GO biological process enrichment analysis and KEGG pathway enrichment analysis were performed for DEGs with the online DAVID Functional Annotation Tool (https://david.ncifcrf.gov/).

### Western blotting

Cells were washed with pre-cooled PBS and proteins were extracted in lysates (Biosharp) supplemented with protease inhibitor cocktail and phosphatase inhibitor cocktail (MedChemExpress). Protein concentration was determined using the Pierce BCA protein reagent kit (Thermo Fisher Scientific). Equal amounts of protein samples were then separated by electrophoresis on a 10% SDS-PAGE gel. The separated proteins were transferred onto a 0.2 μm PVDF membrane (Bio-Rad). The membranes were blocked with 5% nonfat milk (Bio-Rad) at room temperature, then hybridized with primary antibodies overnight at 4L, followed by HRP-conjugated secondary antibodies for 1h at room temperature. The signals were visualized with ECL solution (Bio-Rad) and the protein expression levels were quantified by analyzing the integrated density with Image J software. The antibodies used are listed in Table 3.

### ELISA

The levels of PGD2 in the supernatant of eNK and pbNK cells were measured with a commercial human PGD2 ELISA kit (Elabscience) according to the manufacturer’s instruction.

### Analysis of scRNA-seq

Single-cell sequencing of human endometria was performed by Gene Denovo Biotechnology Co. (Guangzhou). The major cell types were annotated by Seurat and Single R package with a resolution of 0.75. Differential expression analysis (fold change > 2, P < 0.05) was performed to identify genes up-regulated in each subtype. Trajectory analysis of NK subsets was performed by Monocle2. The interactions of NK cells and other cells were investigated by CellPhoneDB package.

### Experimental murine model

Animal experiments were approved by the Ethics Review Board for animal studies of Nanjing Drum Tower Hospital, Affiliated Hospital of Nanjing University Medical School (DWSY-22151278). Eight-week-old C57BL/6 female mice (∼ 20 g) were maintained in SPF conditions with 12 h/12 h cycles of light and darkness, and adapted for one week for the following experiment. Consistent with the late proliferative phase of human menstrual cycle, the model was built at the estrum stage according to the vaginal smear. All mice were anesthetized by inhalation of isoflurane. To establish an IUA-like murine model, the uterus was gently exposed by a mid-abdominal incision, and a 7-gauge needle with rough surface was inserted. And then we scratched the entire uterine wall up and down for about 100 times until the uterus became hyperemic. The abdominal muscularis and skin were carefully sutured, and the uterus was scratched again 5 days later. The mice in the sham-operated group were only given laparotomy. Then the IUA-like mice were randomly divided into 5 groups. For eNK or pbNK cells-treated group, 200 μl supernatant of cells was injected intraperitoneally daily for 10 days. For PGD2-IN-1-treated group, 250 ug/kg PGD2-IN-1 was preinjected 2 h prior to eNK administration. For PGD2-treated group, 200 ug/kg PGD2 was injected intraperitoneally daily for 10 days, and the IUA group received 200 μL blank medium. Mice uterine samples were also harvested at estrus for further testing.

### Statistical analysis

Statistical analysis was performed with GraphPad Prism Version 8.0 (GraphPad Software, Inc). Statistical differences between two groups were analyzed using the unpaired two-tailed Student’s t test and among multiple groups were performed by one-way ANOVA. All data were presented as mean ± standard division (SD) of at least three independent experiments. P value < 0.05 was considered to indicate statistically significant.

## Data availability

The scRNA-seq data used in this study have been deposited in the NCBI Sequence Read Archive (PRJNA730360, PRJNA788201); RNA-seq data of ESCs are available in the NCBI Sequence Read Archive (PRJNA1190156).

## Acknowledgements

This study was supported by research grants from the National Natural Science Foundation of China (82171618, 82371641, 82271653, 82471663), National Key R&D Program of China (2021YFC2701603), Jiangsu Provincial Obstetrics and Gynecology Innovation Center (CXZX202229), Jiangsu Key Laboratory for Molecular Medicine (BM2007208) and Jiangsu Biobank of Clinical Resources (BM2015004).

## Author Contributions

Zhenhua Zhou and Qiao Weng: performing experiments, analyzing data/results, writing original draft, writing - review and editing. Dan Liu and Simin Yao: supporting flow cytometry analysis, analyzing data/results, writing - review and editing. Xueqin Zou, Zihan Zhou and Xier Zhang: supporting animal experiments, writing - review and editing. Huiyan Wang, Hui Zhu and Xiwen Zhang: collecting samples, writing - review and editing. Ling Chen: pathologic analysis, writing - review and editing. Guangfeng Zhao: conceptualization, supervision, writing original draft, writing - review and editing, funding acquisition. Yali Hu: conceptualization; supervision, writing original draft, writing - review and editing, funding acquisition.

## Conflict of Interest

The authors declare no potential conflict of interest.

## Notes

### Competing Interest Statement

The authors have declared no competing interest.

