## Supplementary for "Abnormal eNK cells contribute to endometrial fibrosis in intrauterine adhesions patients"

**Running title: Abnormal eNK cells contribute to endometrial fibrosis**

**This file includes:**

**Supplementary Figures S1-S6**

**Supplementary Figure liagends S1-S6**

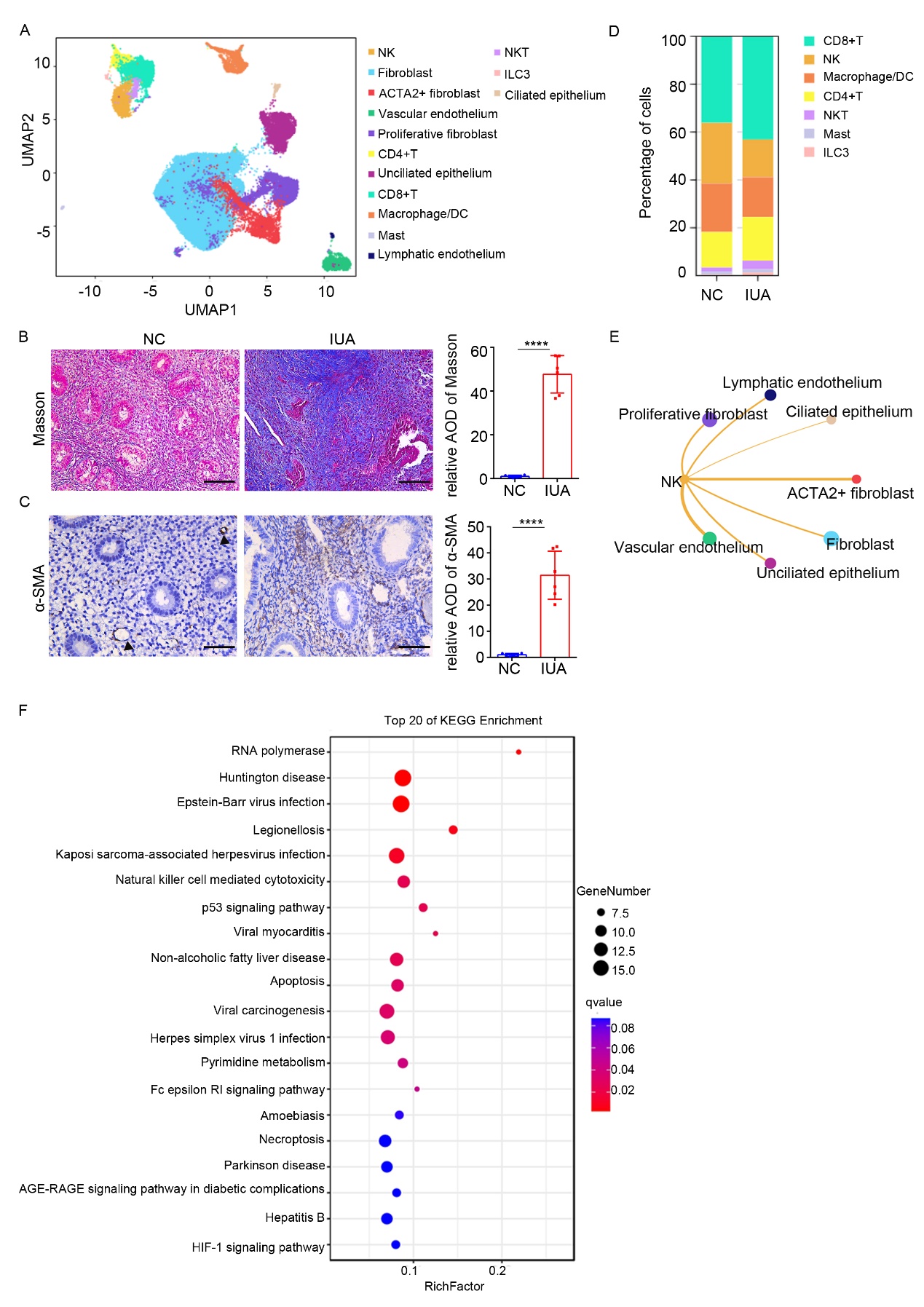

**Supplementary Fig. 1 eNK cells are decreased and myofibroblasts are increased in endometria of IUA patients.**

(A) UMAP plot showing the distribution of 14 major cell clusters in normal and IUA endometria. (B) The Masson staining of endometria in normal controls (n = 6) and IUA patients (n = 6). (C) Immunohistochemical staining of α-SMA in endometria of normal controls (n = 6) and IUA patients (n = 6). Scale bars: 50 μm. (D) The proportion of immune cells (including CD8+T cells, NK cells, Macrophages/DC, CD4+T, NKT, Mast cells and ILC3) in endometria of normal and IUA patients based on scRNA-seq. (E) The CellPhoneDB analysis of the connection between eNK cells (as legend cell) and other non-immune cell types (as receptor cell). (F) Top 20 enriched pathways based on KEGG analysis of the DEGs of eNK cells between normal controls and IUA patients based on scRNA-seq. Error bars, mean ± SD. *****P* < 0.0001.

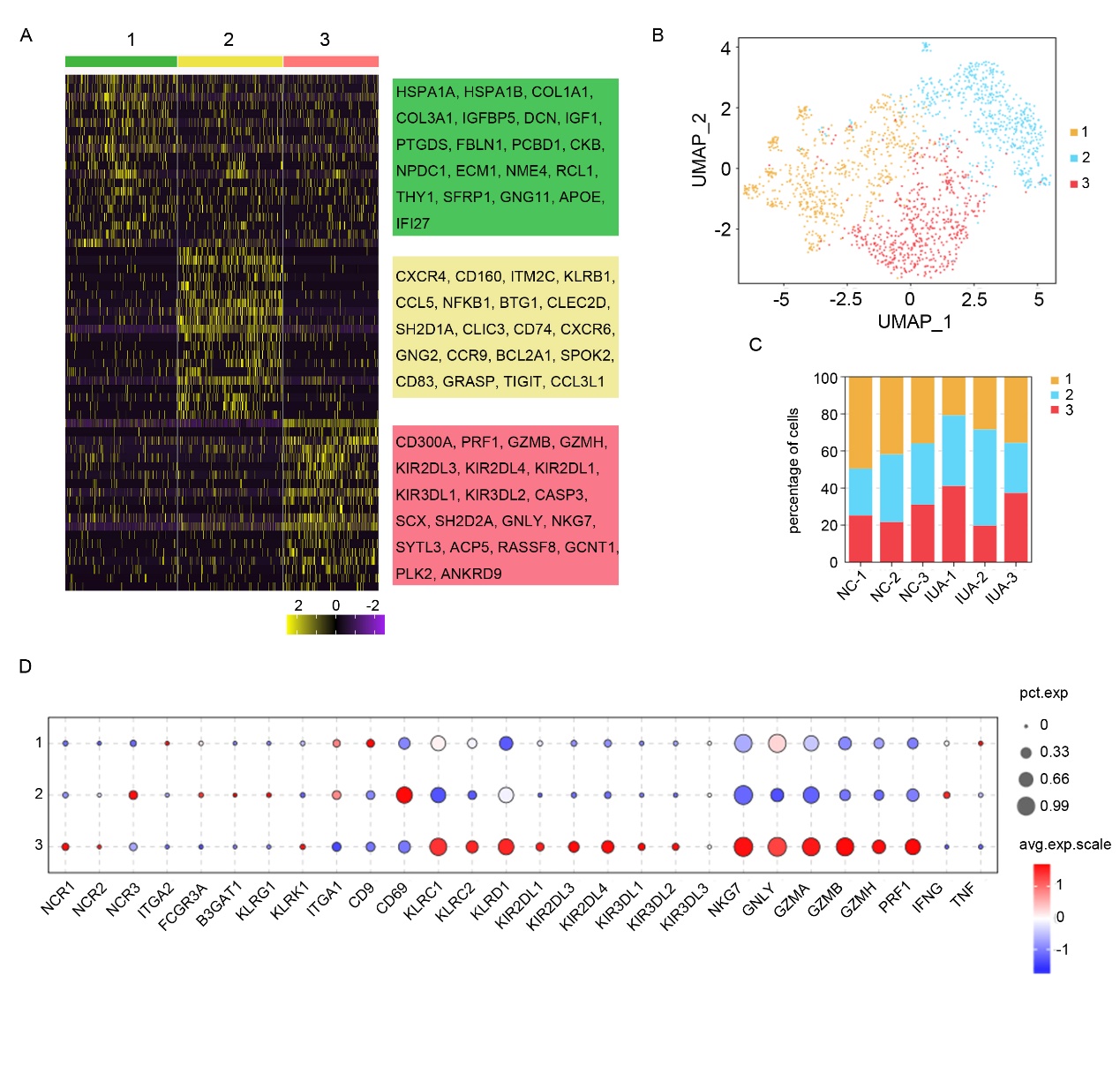

**Supplementary Fig. 2 Sub-clustering of eNK cells.**

(A) Heatmap of top 20 DEGs in each subcluster of eNK cells. (B) UMAP plot of eNK cells. (C) The proportion of each subcluster of eNK cells in normal and IUA patients. (D) Bubble diagram showing the expression of classical surface molecules and effector molecules in eNK cells.

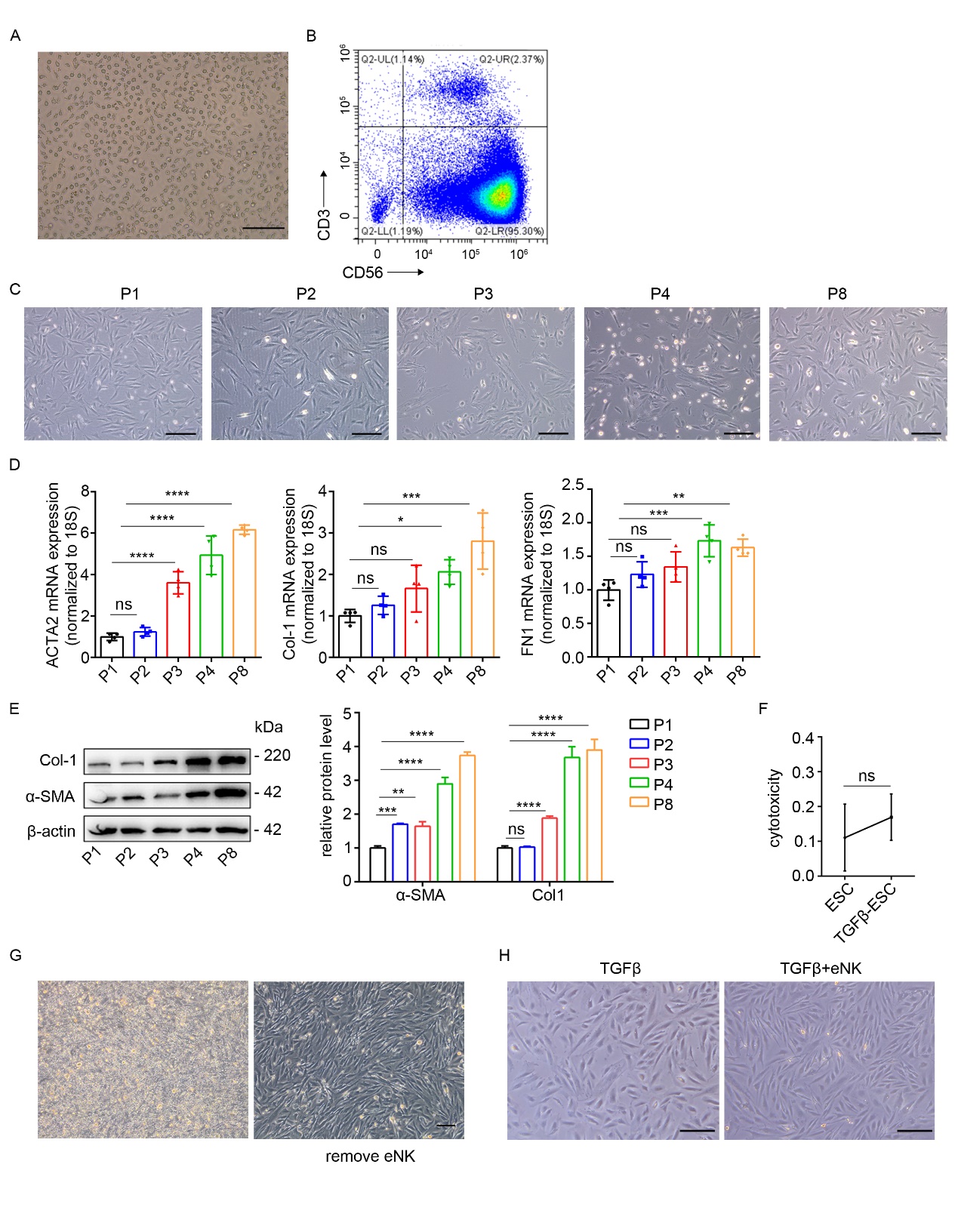

**Supplementary Fig. 3 The isolation of eNK cells and its effect on ESCs and myofibroblasts.**

(A) Representative image of the morphology of eNK cells isolated from endometria in normal controls. Scale bars: 200 μm. (B) Flow cytometric analysis of the isolated eNK cells from normal controls. (C) Representative images of the morphology of ESCs during passage. (P: passage). Scale bars: 200 μm. (D) The mRNA levels of ACTA2, Col-1 and FN1 in ESCs (n = 4) examined by qRT-PCR. (E) Left: The protein levels of α-SMA and Col-1 in ESCs (n = 3) detected by Western Blotting. Right: Relative band intensities analyzed by Image J. (F) The cytotoxicity of eNK cells to ESCs and TGFβ1-induced myofibroblasts (E : T = 10 : 1) analyzed by LDH release assay kit. (G) Left: Representative image of eNK cells co-cultured with TGFβ1-induced myofibroblasts. Right: Representative image of the morphology of TGFβ1-induced myofibroblasts co-cultured with eNK cells. Scale bars: 200 μm. (H) Representative images of the morphology of ESC cells co-cultured with the supernatants of eNK cells. Scale bars: 200 μm. Error bars, mean ± SD. **P* < 0.05, ***P* < 0.01, ****P* < 0.001, *****P* < 0.0001. ns: not significant.

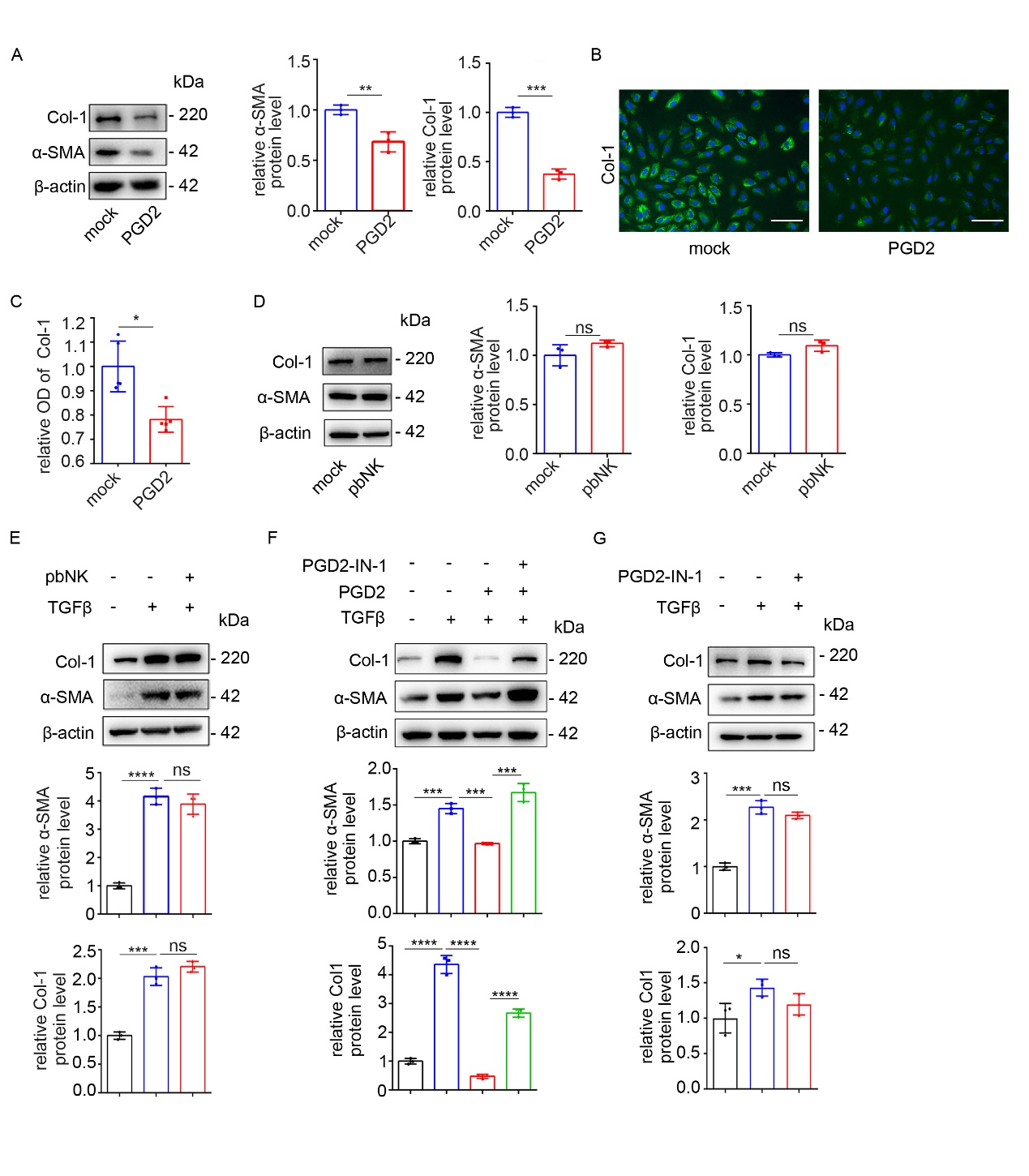

**Supplementary Fig. 4 PGD2 can inhibit the differentiation of ESCs into myofibroblasts.**

(A) Left: The protein levels of α-SMA and Col-1 in ESCs (n = 3) treated with 10 μM PGD2 for 48 h detected by Western Blotting. Right: Relative band intensities analyzed by Image J. (B) Immunofluorescence staining of Col-1 in ESCs (n = 5) treated with the 10 μM PGD2 for 48 h. Scale bars: 100 μm. (C) The relative OD of Col-1 analyzed by Image J. (D) Left: The protein levels of α-SMA and Col-1 in ESCs (n = 3) treated with supernatants of pbNK cells for 48 h detected by Western Blotting. Right: Relative band intensities analyzed by Image J. (E) The ESCs were pre-treated with 10 ng/ml TGFβ1 for 48 h and then incubated with the supernatants of pbNK cells. Top: The protein levels of α-SMA, Col-1 and FN1 in ESCs (n = 3) examined by Western Blotting. Bottom: Relative band intensities analyzed by Image J. (F) The ESCs were pre-treated with 10 ng/ml TGFβ1 and PGD2-IN-1 and then treated with 10 μM PGD2 for 48 h. Top: The protein levels of α-SMA, Col-1 and FN1 in ESCs (n = 3) examined by Western Blotting. Bottom: Relative band intensities analyzed by Image J. (G) The ESCs were pre-treated with 10 ng/ml TGFβ1 for 48 h and incubated with 1 μM PGD2-IN-1 for 24 h. Top: The protein levels of α-SMA, Col-1 and FN1 in ESCs (n = 3) examined by Western Blotting. Bottom: Relative band intensities analyzed by Image J. Error bars, mean ± SD. **P* < 0.05, ***P* < 0.01, ****P* < 0.001, *****P* < 0.0001, ns: not significant.

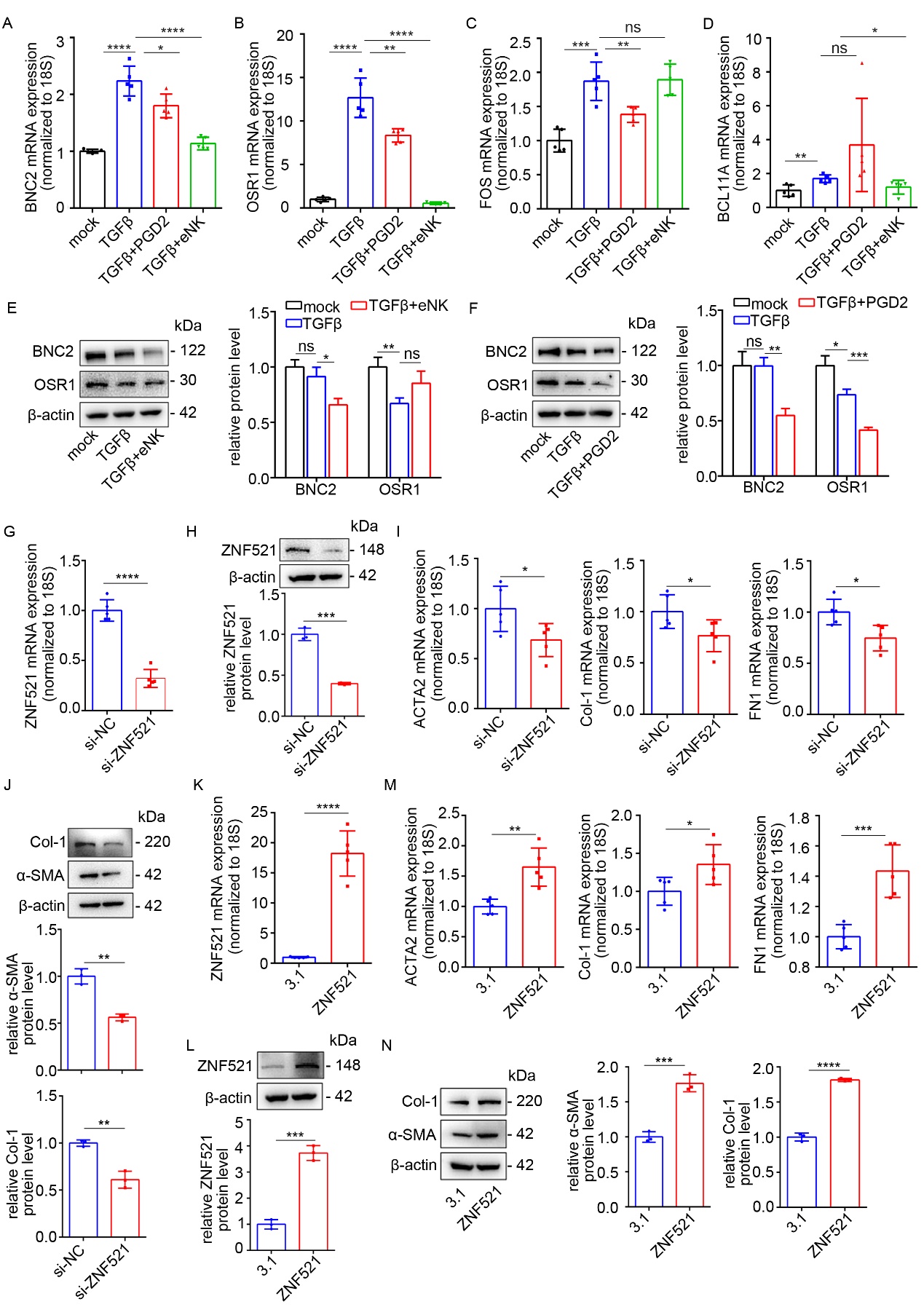

**Supplementary Fig. 5 ZNF521 promotes the differentiation of ESCs into myofibroblasts.**

(A-F) The ESCs were pre-treated with 10 ng/ml TGFβ1 for 48 h and then treated with 10 μM PGD2 or supernatants of eNK cells for 48 h. (A-D) The mRNA levels of BNC2, OSR1, FOS and BCL11A in ESCs (n=5) detected by qRT-PCR. (E-F) Left: The protein levels of BNC2 and OSR1 in ESCs (n = 3). Right: Relative band intensities analyzed by Image J. (G) The mRNA level of ZNF521 in ESCs (n = 5) transfected with si-ZNF521 or si-NC for 48 h detected by qRT-PCR. (H) Top: The protein level of ZNF521 in ESCs (n = 3) transfected with si-ZNF521 for 48 h detected by Western Blotting. Bottom: Relative band intensities analyzed by Image J. (I) The mRNA levels of ACTA2, Col-1 and FN1 in ESCs (n = 5) transfected with si-NC or si-ZNF521 for 48 h examined by qRT-PCR. (J) Top: The protein levels of α-SMA and Col-1 in ESCs (n = 3) transfected with si-NC or si-ZNF521 for 48 h detected by Western Blotting. Bottom: Relative band intensities analyzed by Image J. (K-L) The mRNA (n = 5) and protein level (n = 3) of ZNF521 in ESCs transfected with ZNF521 plasmid for 48 h. (M) The mRNA levels of ACTA2, Col-1 and FN1 in ESCs (n = 5) transfected with ZNF521 plasmid for 48 h detected by qRT-PCR. (N) The protein levels of α-SMA and Col-1 in ESCs (n = 3) transfected with ZNF521 plasmid for 48 h detected by Western Blotting. Error bars, mean ± SD. **P* < 0.05, ***P* < 0.01, ****P* < 0.001, *****P* < 0.0001, ns: not significant.

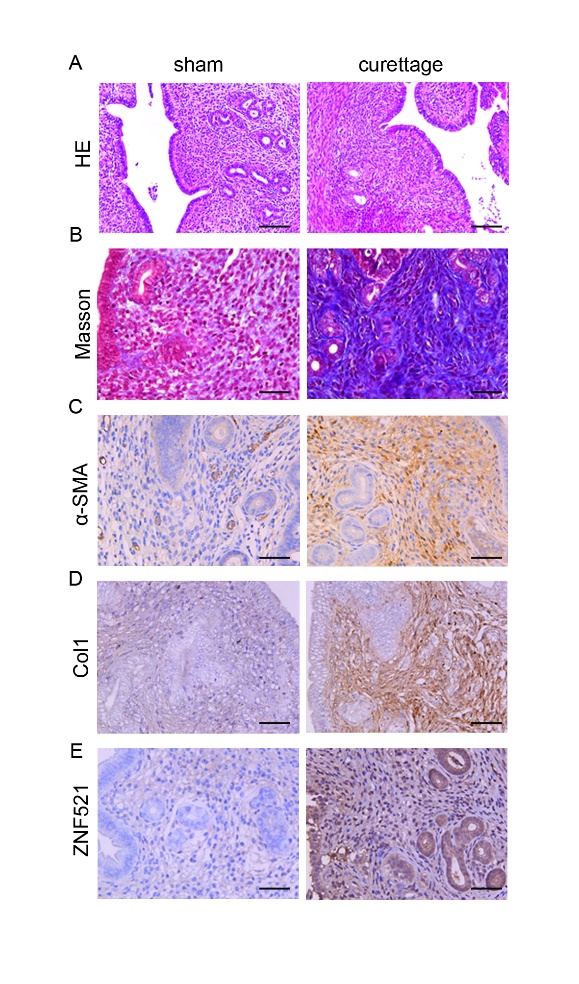

**Supplementary Fig. 6 The establishment of the IUA murine model.**

(A) HE staining in uterus of mice with sham operation and IUA model. Scale bars: 100 μm. (B) Masson staining of uterus of mice with sham operation and IUA model. Scale bars: 50 μm. (C-E) Immunohistochemical staining of α-SMA, Col-1 and ZNF521 in uterus of mice with sham operation and IUA model. Scale bars: 50 μm.
